# “The anti-apoptotic effect of *Lonomia obliqua* hemolymph is associated with the mitochondria pathway”

**DOI:** 10.1101/2022.06.02.494522

**Authors:** Ronaldo Z. Mendonça, Luciana Moreira Martins

## Abstract

The apoptosis death is a very important factors in production processes that limited the industrial production of some proteins of economic interest. However, one of the forms to increase the cellular productivity would be inhibit or attenuate the cellular death. Recently we have demonstrated the presence of a potent anti-apoptotic protein in *Lonomia obliqua* hemolymph which extends the cell culture viability through apoptosis prevention. By the other side, has been reported that mitochondria have one important action in the apoptosis control process, being that mitochondria membrane permeabilization (MMP) can be an important stage in this process. MMP associated or not with the loss of the electrochemical potential of the mitochondria and alteration of the matrix is responsible for the inter membranes protein release (e.g. cytochrome c, the AIF, etc) of cytosol. The result obtained showed that the addition of a protein from *Lonomia obliqua* hemolymph in the culture lead to a prolongation of the cellular life (3-4 days) and the cells leading a high electrochemical potential of the mitochondria. This protein can has its action in mitochondria membrane, avoiding the loss of the membrane permeability and the Cytochrome-C release. As positive control, apoptosis death in these cultures was induced by 50 μm of t-BHP or 600 μm of H_2_0_2_. The presence of apoptosis was characterized by flow citometry, microscopy electronic and agarose gel electrophoresis. The potential electrochemical of the mitochondria was determined by JC-1, Hoechst 33324 and DIOC6. Cytochrome C was identified in cytosol by an anti-cytochrome antibody.

## Introduction

Optimization of animal cell culture processes is essential for economical production of biopharmaceutical products, such as recombinant proteins, viruses and cells. Cell death in bioreactors represents a major problem in cell culture technology by decreasing the global productivity yield. Programmed cell death is directly related to the cells death in bioreactor, being 2 mechanisms of death can be observed; apoptosis and autophagy. The programed cell death (apoptosis) mechanism is activated by various external and internal factors as heat, irradiation nutrient depletion, shear stress, hypoxia, toxin accumulation, as well as a cell response to viral infections, (Kazek et al., 2020; Cao et al., 2021; Nury et al., 2021; Wang et al., 2021; Wrońska et al 2022, Mendonça et al. 2008; Arden et al. 2007; Mendonça et al. 2002; Meneses-Acosta et al. 2001;). These factor act directly on the membrane of cells or on mitochondria (Bedoui et al., 2020), being acting on a caspase activation cascade (Bock and Tait, 2020; Kesavardhana et al., 2020). Since these factors implicated on the lost of cell viability, due to cell death by apoptosis, the decreasing apoptosis in cell culture is an attractive strategy to improve global productivity yields (Butler 2005). For that, two main strategies can be followed: manipulation of the external environment or genetic manipulation of the internal cellular biochemistry. The medium culture can be supplemented with various components for prolongation of the viability cell. These components can be anti-apoptotic agents, nutrients or selected agents from serum. Caspases inhibitors also can enhance cell culture viabilities and protein titer (Arden et al. 2007; Sauerwald et al. 2003). Anti-oxidants were also described as factors to improve cell viability, N-acetylcysteine increases mammalian cell lifetimes upon Sindbis virus vector infection (Mastrangelo et al. 1999) and catalase can improve recombinant protein production in baculovirus-insect cell system by preventing cell death (Vieira et al. 2006). We have showed that supplementation of insect cell culture with *Lonomia obliqua* hemolymph could extend culture longevity, avoiding apoptosis death (Maranga et al. 2003). Earlier we have identified, isolated and characterized this protein from *Lonomia obliqua* hemolymph (with about 51 kDa), presenting a potent anti-apoptotic effect. Addition of this purified protein to Sf-9 cell cultures allowed the prevention of apoptosis induced by nutrient depletion, as well as by actinomycin D. This protein showed action as in mammalian cell as well in insect cel showing a broad action range (Souza et al. 2005).

After this, when this protein was added to insect infected cell culture, was observed an improvement of recombinant rabies virus glycoprotein (rRVGP), expressed by Drosophila melanogaster Schneider 2 (S2) (Mendonça et al. 2008); and improvement of recombinante bovine rotavirus glycoprotein Vp2, Vp6 and Vp7, expressed by baculovirus (Vieira et al. 2010). Recently, we have observed that the addition of this protein to cell culture also leads to an increase in baculovirus production. This increase in viral production could be related to the increase in the production of recombinant proteins by baculovirus (Souza et al. 2015). So, in this work, we seek to identify the place of action of this protein. Mitochondrial membrane permeabilization (MMP) can be a rate-limiting step of apoptotic just as of necrotic cell death. MMP associated or not to the loss of mitochondrial electrochemical potential (ΔΨm) and matrix swelling, is responsible for the release of intermembrane proteins (e.g. cytochrome Cc, AIF, etc) into the cytosol. As a result of this release, apoptogenic proteases (caspases) and DNases are activated and the death process becomes irreversible. In addition, MMP can cause depletion of antioxidant enzymes, reactive oxygen species generation and cessation of ATP synthesis leading to an oxido-reductive and bioenergetic catastrophe (Kroemer et al. 2007; Ferri et al. 2000; Ferri and Kroemer 2001). Therefore studying the implication of mitochondria on the apoptosis inhibition by *Lonomia obliqua*, hemolymph factors is highly important to acquire a better knowledge about their mechanisms of action.

The present work aims to demonstrate the anti-apoptotic ability of hemolymph factors in anchorage-dependent HEK-293 cells cultures. The involvement of *Lonomia obliqua* hemolymph in the mitochondrial control of apoptosis is disclosed.

## Materials and methods

### Cells and media

Anchorage-dependent HEK-293 cells, purchased from ATCC (ATCC-CRL-1573), were routinely cultured in Minimum Essential Medium (MEM) supplemented with 5% (v/v) heat-inactivated (56°C, 30 min) Fetal Bovine Serum (FBS), 2mM of glutamine, using an humidified atmosphere of 5% CO_2_ in air at 37°C.

### Total soluble protein quantification

The total soluble protein content of the whole hemolymph and purified protein fraction were quantified using the Bradford method (Bradford 1976) using bovine serum albumin as a standard.

### Hemolymph

Hemolymph of *Lonomia obliqua* was collected from sixth-instar larvae after setae cut off. The collected hemolymph was centrifuged by 1.000g/10 min; the supernatant was filtered with 0.2μm membrane filter, inactivated by heat (60°C/30min) and storage at 4°C. This material was used for supplementation the medium (1%) at start of the culture.

### Hemolymph purification by chromatography

After centrifuged and filtered, 1 ml of hemolymph was loaded on a Superdex^™^ 75 10/300 GL (Amersham Pharmacia Biotech) column at rate of 0.5 ml/min and eluted with sodium phosphate buffer. The eluated was harvested in fractions of 0.5 ml and monitored at 280 nm. Active fractions from Superdex^™^ 75 10/300 GL column were loaded at an ion change column (Resource q). The chromatography was performed at an AKTA purifier chromatograph (Amersham Pharmacia Biotech). The purified fractions were applied at SDS-PAGE electrophoresis for analysis.

### Chemical inductor of apoptosis

Apoptosis was trigged by oxidative stress induced with 50, 75 or 100μM of t-BHP (Tert-butylhydroperoxide) (Sigma) or H_2_0_2_, in the concentrations of 400, 600, 800 or 1000 μM.

### Fluorochromes used to determination of cellular death

Propidium iodide (1μg/ml) and Hoechst 33342 (2 μM) was used as indicator of cellular death. The propidium was used as a fluorocrome that penetrates in all of the cells that lost the permeability emitting in red. Hoechst is a fluorochrome that stain cell nucleus allowing the visualization of morphologic changes of nuclei in apoptosis cell death process.

### Fluorescent microscopy

HEK-293 cells were cultured in 13 mm-diameter cover slips. Eighteen hours later, cells were treated with tert-butylhydroperoxide (Sigma) and 4 hours after cells were stained with Hoechst 33342 (2μM, Sigma), followed by fluorescence microscopic assessment of apoptotic nuclei. Cells were observed on a Leica DMRB microscope using a filter cube presenting UV excitation range with a band pass of 340-380 nm of wave length.

### Flow cytometry

Samples of 0.5 ml were collected from the cell culture at different times. Apoptosis-associated changes were assessed by cytofluorometry on a BD FACSCalibur^™^ 4 colors (Becton Dickinson), while gating the forward and the side scatters on viable cells, using several fluorochromes: 3,3’ dihexyloxacarbocyanine iodide (DiOC6(3), 20 μM) for ΔΨm quantification, propidium iodide (PI, 1 μg/ml) for the determination of cell viability. The acquisition and analysis of the results was performed with the software Cell Quest (Becton Dickinson).

### Mitochondrial membrane potential measure

JC-1 (5,5’,6,6’-tetrachloro-1,1’,3,3’-tetraethylbenzimidazolyl-Carbocyanine iodide) (2μm) and DIOC6 (3) (3,3’dihexyloxacarbocyanine iodide), (20μM) were used as mitochondrial membrane potential indicator.

### Immunofluorescence for cytochrome C detection

For identification of cytochrome C was used a monoclonal antibody anti-cytochrome C and a second IgG anti-mouse antibody conjugated with alkaline phosphatase.

### Measurement of cell viability

Culture samples were obtained daily and cell concentration was measured using a hemocytometer. Cell viability was determined by trypan blue exclusion test under light microscopy.

### Participation of the mitochondria in the apoptosis death process

The participation of the mitochondria during the cellular death by induced apoptosis and the protector effect of the hemolymph was determined by the potential of the mitochondrial membrane. To this, apoptosis death in HeK-293 cells was induced by t-BHP or H_2_0_2_. The identification of the potential of the mitochondrial membrane in the cells cultured was accomplished by FACs (suspension cells) after cells stains with DIOC_6_ (3) or in fluorescence microscope (adherent cells) after cell stain with JC-1 / Hoechst 33324. The pro-apoptotic factors (cytochrome C) determination liberated in the cytosol of the HEK-293 cell mitochondrial, after apoptosis induction, was performed by an anti-cytochrome C antibody, as described below.

### Determination of the potential of the mitochondrial membrane in HEK - 293 cell after JC-1 stain

For the determination of the protector effect of hemolymph against apoptosis induced (t-BHP or H_2_0_2_), cells HEK-293 were cultivated in suspension or adherent to micro plates. After semi confluence, cells were treated with 1% (v/v) of total hemolymph (Hb) or with purified fraction (Frp) showing anti-apoptotic activity. After one hour of contact cells were treated with 50, 75 or 100μM of t-BHP or with 400, 600, 800 or 1.000μM of H_2_0_2_. The cells cultures were maintained at CO2 incubation by 4 hours and after this period cells were washed with PBS and treated with Hoechst 33324 and the fluorochrome JC-1 (2μM).

The determination of the cells potential of the mitochondrial membrane was performed by FACs (FACSort Becton Dickinson (Air laser, to 488 nm (excitement) and 620 nm

(emission), for cells in suspension or in fluorescence microscope for adherent cells). Cells showing mitochondria with low and high potential of the mitochondrial membrane are stained in green and red respectively. Normal cells present homogeneous nucleus stained in blue (Hoechst) while apoptotic cells present the nucleus fragmented.

### Identification of Cytochrome C in the cytosol of the mitochondria by immunofluorescence

To the determination of the protector effect of the hemolymph in the apoptosis induced by t-BHP, cells HEK-293 were cultivated in micro plates of 12 wells. After semi confluence, the cells were treated with 1% (v/v) of total hemolymph (Hb) or it’s purified fraction (Frp). After contact of one hour the cells were treated with 50 or 75μM of t-BHP. The cultures were incubated by 4 hours and after this period, cells were washed with PBS and fastened with paraformaldeide (4%) and picric acid (0,18%) and incubated by 45 minutes. The cells were washed with PBS, treated with BSA (3%) and incubated by 30 minutes. The cells were then washed with PBS and marked with the cytosol monoclonal antibody. The reaction was revealed with an anti-mouse IgG antibody conjugated with alkaline fosfatase by 75 minutes being after this added 2 μM of hoechst. The material was observed in a fluorescence microscope (microscope Leica DMRB).

## Results

### Hemolymph purification and apoptosis identification

The protein antiapoptotic was purified in two steps of chromatography (Souza et al. 2005). The results are shown in **Figure 1**.

**Figure 1.**
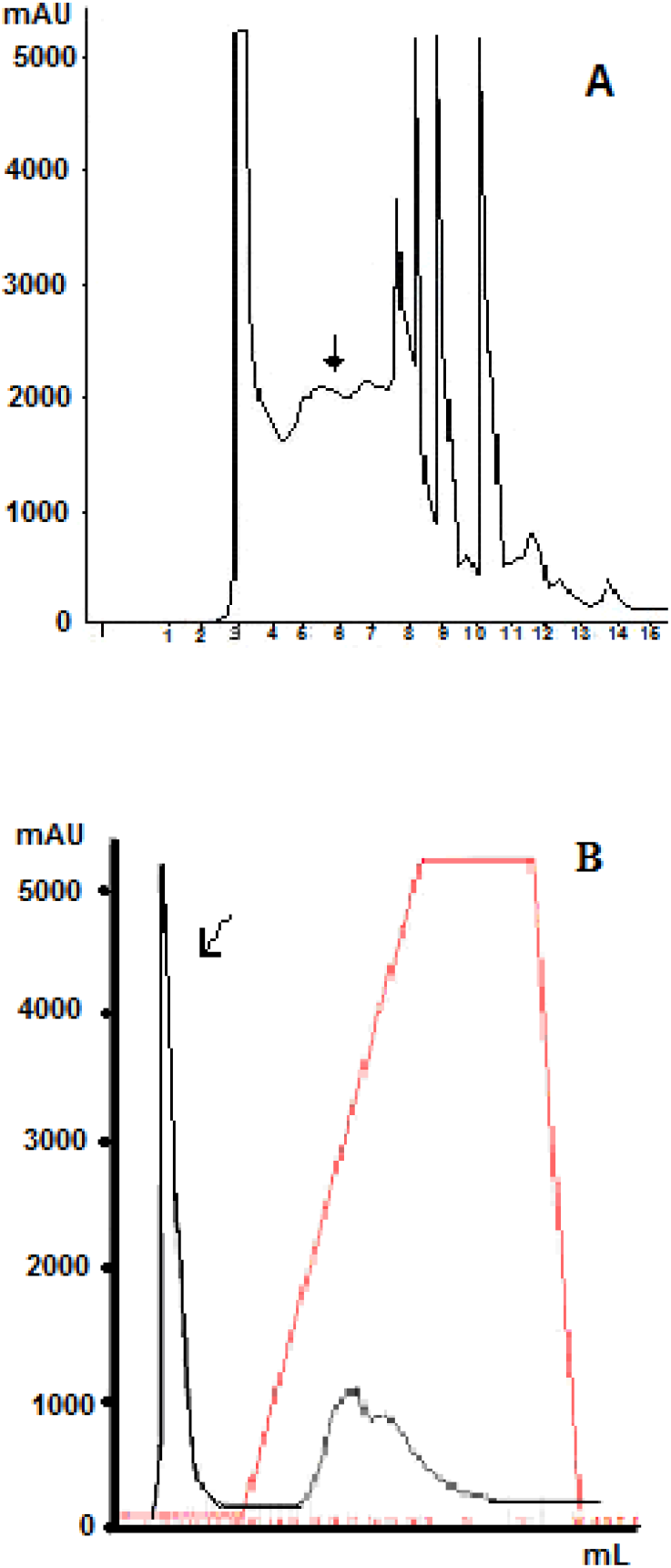
Purification of total hemolymph of *Lonomia obliqua* and its subfractions. Total hemolymph was initially purified by gel filtration in a Superdex G-75 column (Fig 1a). Fractions were tested for antiapoptotic activity. The chromatography fraction with activity (arrows in Figure 1a) was applied on an ion exchange column (Resource Q). The chromatography fraction showing antiapoptotic effect is presented with a arrows in Figure 1b.

The whole and purified fraction was then applied to SDS-PAGE gel electrophoresis. The purified protein against death effect appeared as one band with molecular weights of approximately 51 kDa (**Figure 2**).

**Figure 2.**
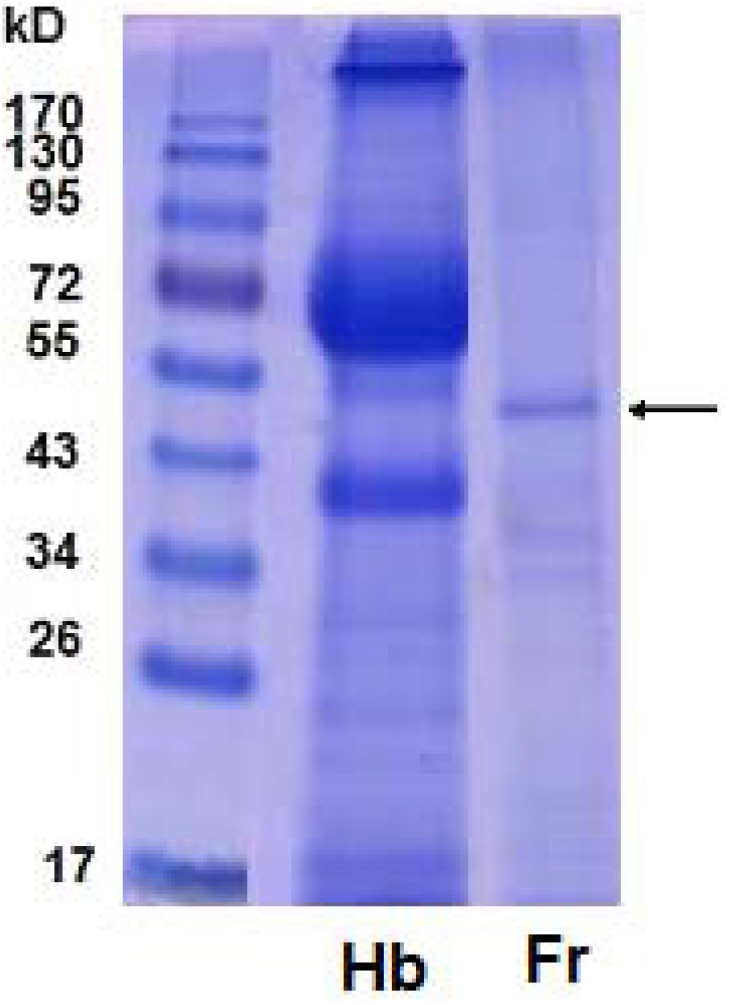
SDS-PAGE of total hemolymph (Hb) and the purified antiapoptotic protein (Fr) analyzed on a 12% gel.

In order to evaluate the effect of hemolymph supplementation in cell death by apoptosis process, cultures were incubated with 1% v/v of whole hemolymph (30.0 mg/ml) or purified protein fraction (1.0 mg/ml). The final concentration in culture was 300 μg/ml and 20 μg/ml respectively. The results obtained in hemolymph supplemented cultures were compared with those cultures performed without supplement addition. The results showed that hemolymph supplementation (whole and purified fractions) had a strong positive protection effect in cell culture, suggesting the presence of a potential anti-apoptotic in hemolymph of *Lonomia obliqua* **(Figure 3).**

**Figure 3.**
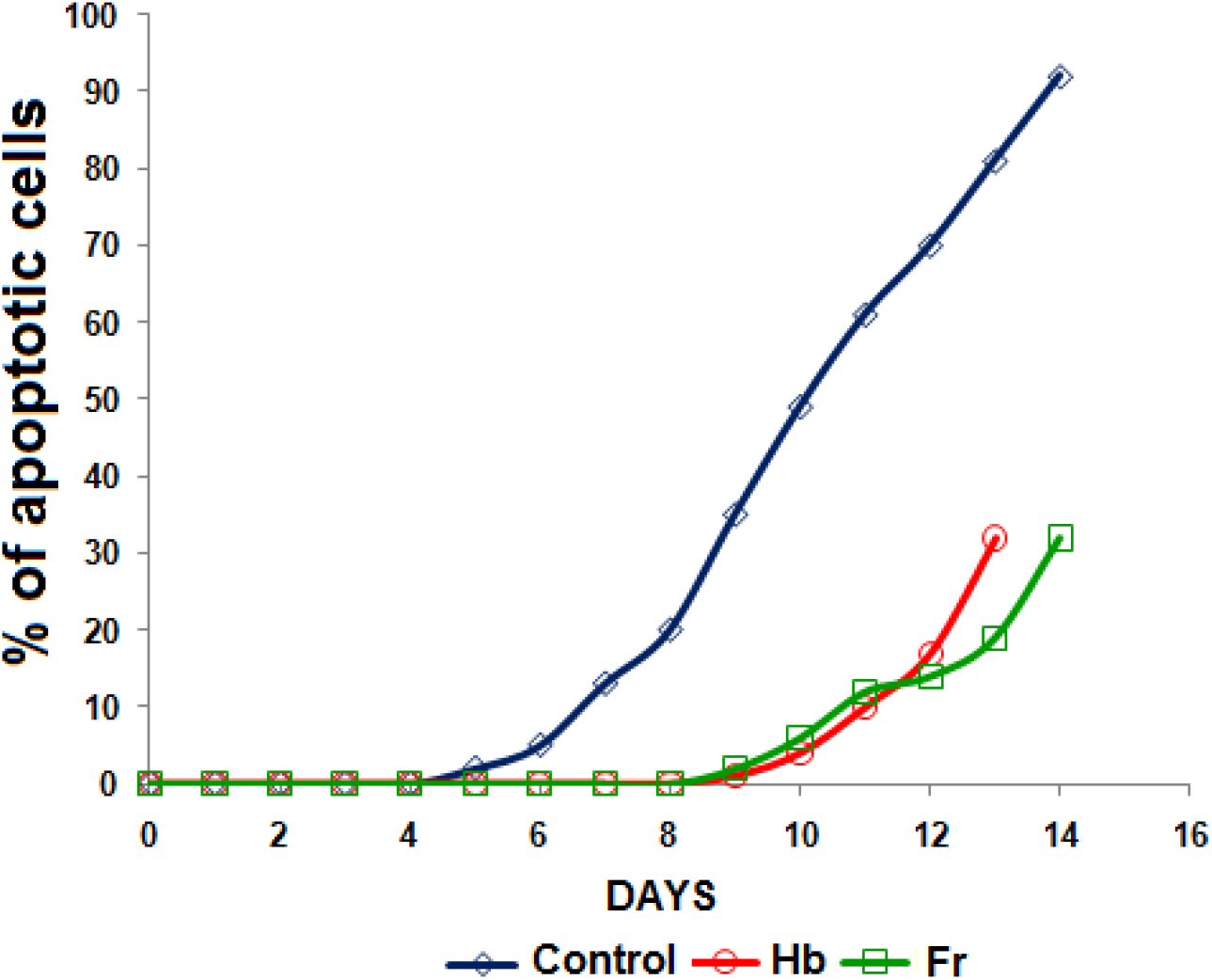
Percentage of apoptotic cells in HEK-293 cultures treated with 1% v/v of total (red) or purified fraction of hemolymph (green). Culture samples were obtained daily and cellular death was determined after propidium iodide (1μg/ml) and Hoechst 33342 (2 μM) stain.

### Determination of the potential of the mitochondrial membrane of HEK-293 cell after induction by t-BHP and H_2_0_2_

#### Effect of different amounts of H_2_0_2_ in the mortality and in the potential of the mitochondrial membrane of HEK-293 cells

Aiming to verify the action of the anti-apoptotic protein in potential of the mitochondrial membrane of HEK-293 cells after induction by t-BHP and H_2_0_2_, HEK-293 cells were previously treated with 1% (v /v) whole or a purified fraction of hemolymph. After 1 hour, cell death was induced with H_2_0_2_ (600, 800 or 1.000μM) overnight. After this period the cultures were labelled with 1 μg/ml PI and 20μM of DIOC6(3). After 20 minutes, the samples were applied to a flow cytometer and the potential of the mitochondrial membrane was determined. As can be seen in **Figure 4**, there is an increase in the loss of potential of the mitochondrial membrane after H_2_0_2_ addition. By the other hand, both, total (Hb) and purified hemolymph fraction (Frp) were able to inhibit cell death, keeping potential of the mitochondrial membrane higher than that observed in control culture. The loss of potential of the mitochondrial membrane (and consequently cell death) was proportional to the increase in the inductor amount. The total hemolymph was able to inhibit virtually almost all cell death, even with 1.000 μm H_2_0_2_. In this condition, the number of dead cells in control was 62%, while that, in the treated hemolymph culture, the amount of cell death did not exceed 4.75%. Even the fraction was able to inhibit the cell death but in lower amount (37%). HEK-293 cells were very sensitive to incubation under the experimental conditions (overnight). In all experiments it was observed a loss of cell viability in control cultures in this period (26.88%), but loss was not observed in cultures incubated with total hemolymph (2.65%). These data suggest that hemolymph also protect the cells of control against initial loss of viability.

**Figure 4.**
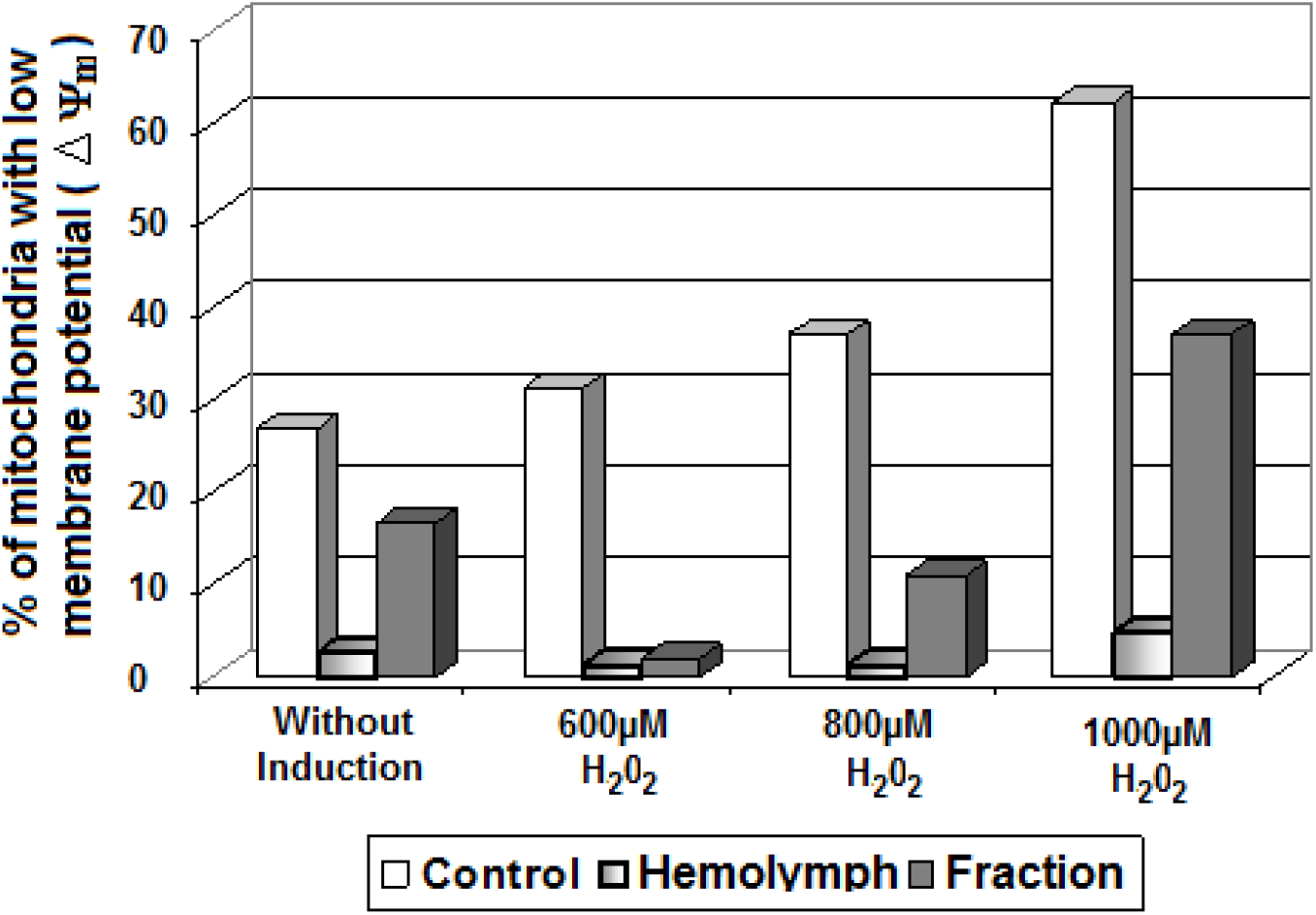
Effect of hemolymph in the protection of the cellular death by H_2_0_2_. HEK-293 cells were previously treated or not with 1% (v/v) of total or purified hemolymph. After this period the cellular death was induced with 600, 800 or 1000μM of H_2_0_2_ overnight. After this period the cultures were stained with 1μg/ml of PI and 20μM of Dioc(6)3. After 20 minutes of contact the samples were applied at FACS and the death number cells were determined.

To verify the protective effect in the potential of the mitochondrial membrane of cells, HEK-293 cells treated with 1% (v /v) of total or purified fractions of hemolymph. After 1 hour, cell’s death induced with 400 or 800 μM of H_2_0_2_, for 4 hours. After this period the cultures were labelled with DIOC6 (3) and the fluorescence was determined. At **Figure 5** is observed the % of fluorescent cells showing high mitochondrial membrane potential. As can be observed, the hemolymph and it purified fraction were able to remain the membrane potential even after treatment of cells with 800 μM of H_2_0_2_. Moreover, the control of the culture incubated with total hemolymph or it fraction, shown higher viability when compared with the control of the culture without this treatment.

**Figure 5.**
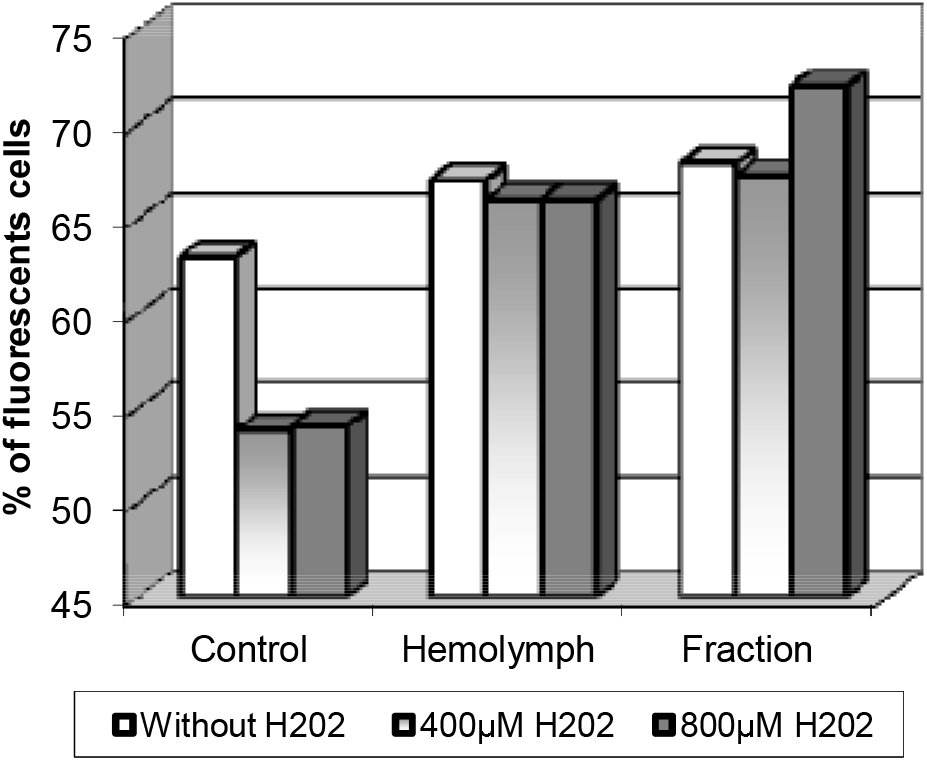
Hemolymph effect in the membrane potential of cells HEK-293. The HEK-293 cells were previously treated or not with 1% (v/v) of total or purified hemolymph. After this period the cellular death of the cells was induced by 4 hours with 400 or 800μ M of H_2_0_2_. After this period the cultures were stained with 1μg/ml of PI and 20μM of DIOC6(3). After 20 minutes of contact, the number of cells showing high fluorescence was determined.

#### Effect of different amounts of t-BHP in the potential of the mitochondrial membrane of HEK-293

At **Figure 6** is showed the effect of different concentrations of t-BHP on membrane potential of mitochondria in HEK-293 cells (four hours of induction). As can be seen there is a progressive increase in the loss of membrane potential of mitochondria according to increasing concentrations of t-BHP. To verify the protective hemolymph effect in the potential of the mitochondrial membrane, HEK-293 cells treated with 1% (v /v) of total hemolymph or it purified fraction. After 1 hour, cell death induced with 50, 75 or 100 μM of t-BHP, for 4 hours. After this period, the cultures were labelled with DIOC6 (3) and fluorescence determined by flow cytometry. At **Figure 7** is observed the percentage of fluorescent cells showing high mitochondrial membrane potential. As can be observed, total and hemolymph fraction were able to inhibit cell death, maintaining the potential of the mitochondrial membrane higher than that observed in control culture.

**Figure 6.**
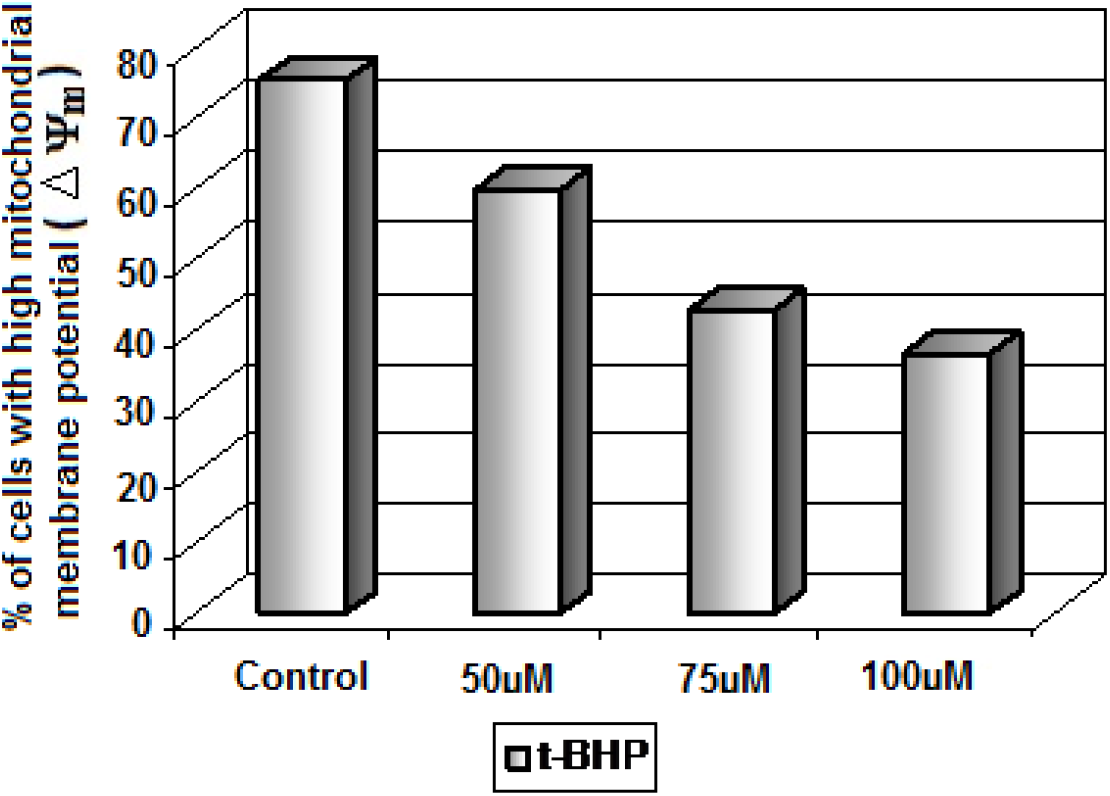
Hemolymph effect in the membrane potential of cells HEK-293. The cellular death was induced by 4 hours with 50, 75 or 100μM of t-BHP. After this period the cultures were stained with 1μg/ml of PI and 20μM of DIOC6(3). After 20 minutes of contact the samples were applied to a FACs and the membrane potential was determined.

**Figure 7.**
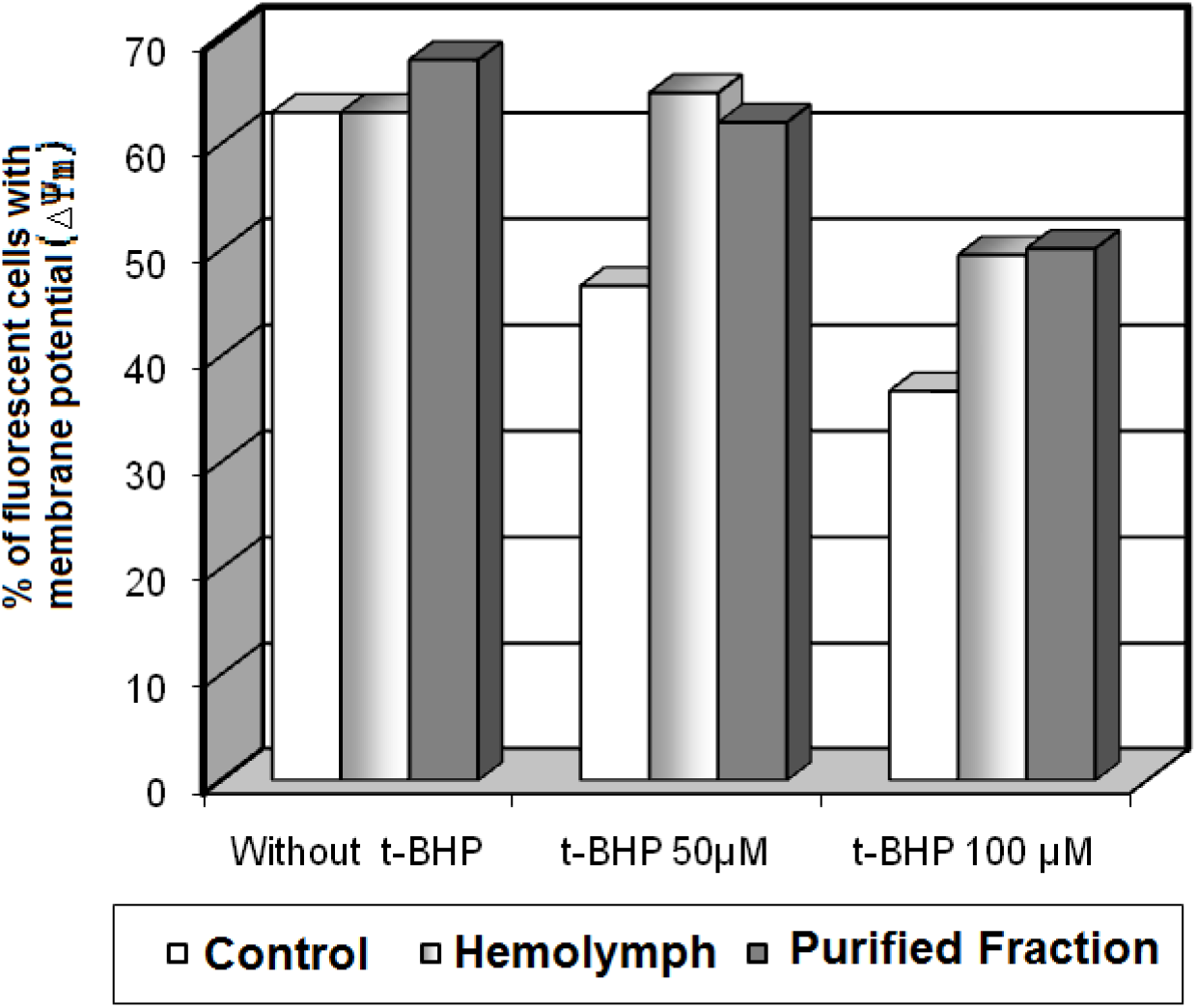
Hemolymph effect in the membrane potential of cells HEK-293. The cells previously treated or not with 1% (v/v) of total or purified hemolymph. After this period the cellular death of the cells was induced by 4 hours with 50 or 100μM of t-BHP. After this period the cultures were stained with 1μg/ml of PI and 20μM of DIOC6(3). After 20 minutes of contact the number of cells showing high fluorescence was determined.

#### Determination of potential of the mitochondrial membrane in the HEK-293 using JC-1 / Hoechst 33324 staining after cell death induction by t-BHP and H_2_0_2_

In the **Figure 8** is showed a comparison between the protector effect of hemolymph and the purified fraction in the HEK-293 cells culture treated with 50 μM of t-BHP or 600 μM of H_2_0_2_. To this, HEK-293 cells grew adhered on the glass slides and treated with total hemolymph or purified fraction 1% (v/v). After 1 hour, cell’s death was induced with 50 μM of t-BHP or with 600 μM of H_2_0_2_, and maintained overnight. Then, the cells were marked with 1μg/ml of PI and 20μM of JC-1 and Hoechst. The potential of the mitochondrial membrane (A) and t he number of nucleous (B) with apoptotic morphology respectively were determined by fluorescent microscope. As can be observed, there is a good correspondence between the potential of the mitochondrial membrane and the number of nuclei with apoptotic morphology.

**Figure 8.**
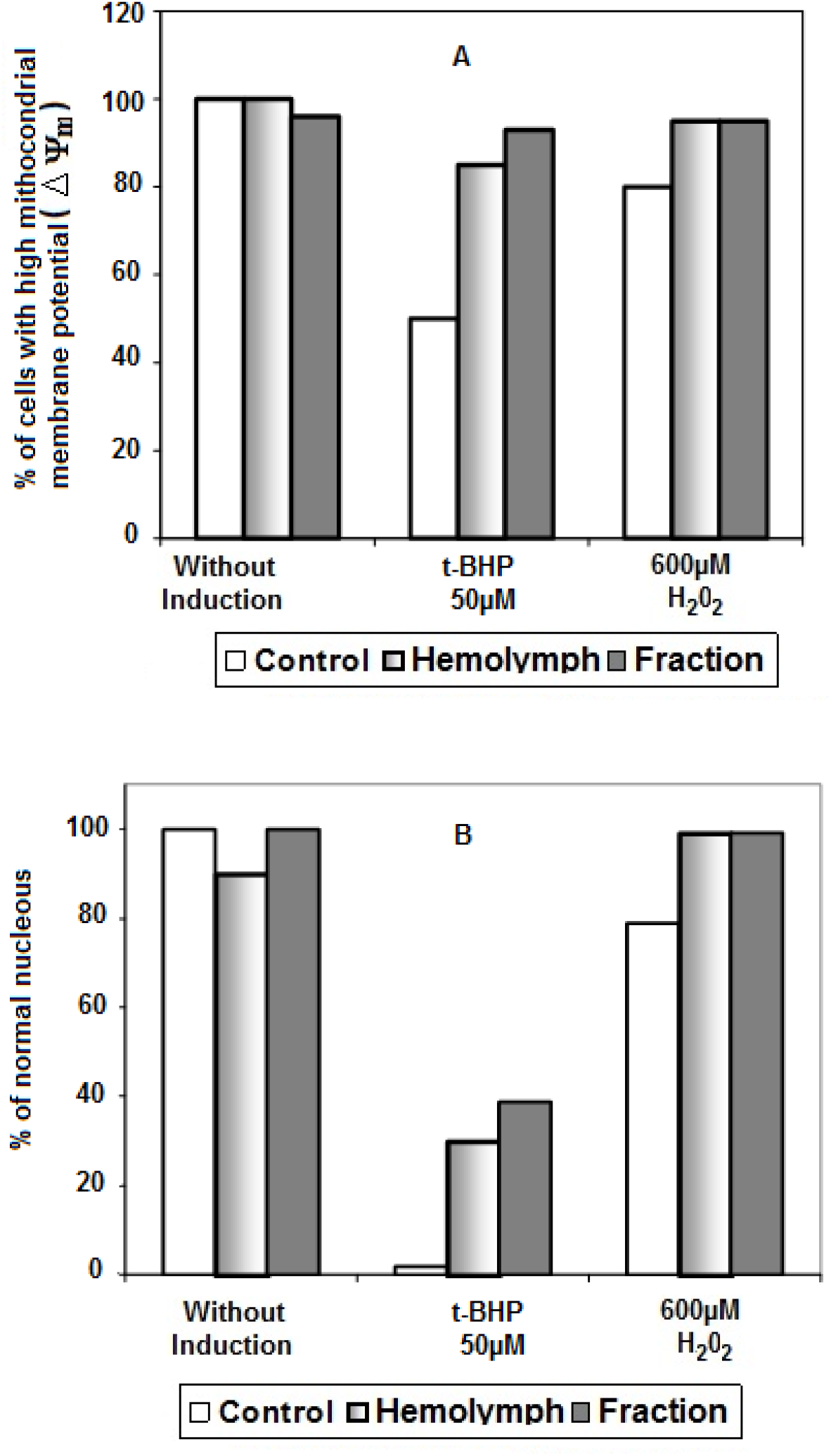
Effect of the hemolymph in the protection of the cellular death by apoptosis chemical inductor. HEK-293 cells previously treated or not with 1% (v/v) of total or purified hemolymph. After this period the cellular death was induced with 50μM of t-BHP or 600μM of H_2_0_2_ overnight. After this period the cultures were marked with 1μg/ml of PI and 20μM of JC-1 and Hoechst. After 20 minutes of contact the samples were applied to a FACs and the membrane potential was determine A). The number of cells with fragmented nucleus was determined by fluorescence microscope.

As showed at **Figure 9**, also there is a good correspondence between the results obtained from both dyes (t-BHP (1st column) and H_2_0_2_ (2nd column) after JC-1 and Hoechst stain. In **Figures 9 A and B** and **E, F** are shown viable HEK-293 cells showing high potential of the mitochondrial membrane. After **t-BHP** or H_2_0_2_ induction, almost all cells were stained green (**Figure 9 C and D**). It was correlated with percentage of fragmented cell nucleus after Hoechst stain showing that the loss of potential of the mitochondrial membrane is related with the cell death and DNA fragmentation **(Figure 9 G and H).** On the other hand, when total hemolymph or its fraction was added, this effect was not observed **(Figure 9 K).** The **Figures 9 I and J** corresponds to the overlap of staining with Hoechst and JC-1. In the **Figure 9 K and L** are presented two pictures with overlap of the two markers, the first cell were apoptosis induced and treated with hemolymph and the second the cells were apoptosis induced but without treatment with hemolymph. These figures are representative of all experiment performed with the cultures were death were inducted by t-BHP or H_2_0_2_ and treated or not with hemolymph. The results obtained in cultures treated with total or fraction hemolymph were identical to the normal controls without induction.

**Figure 9.**
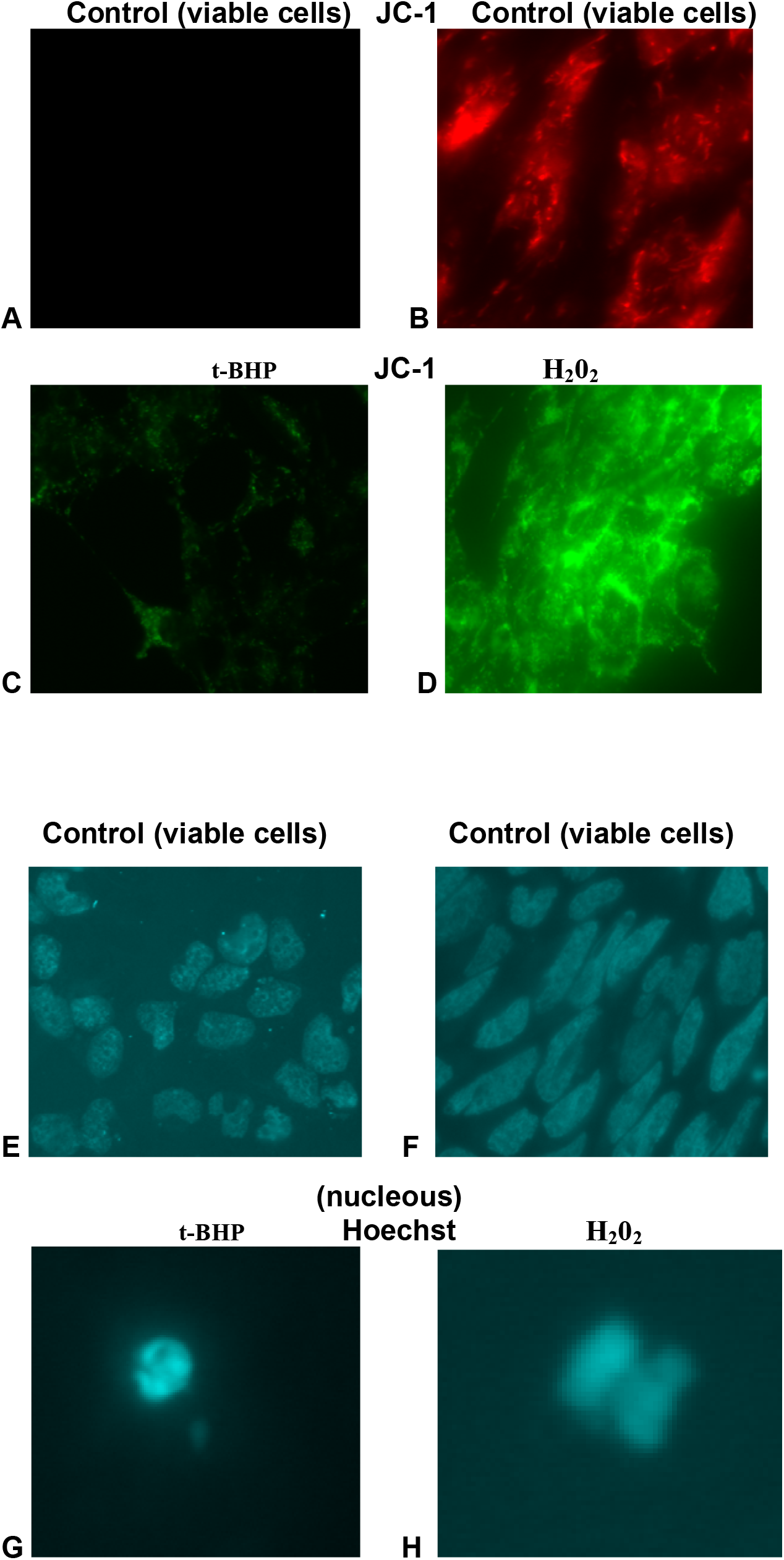
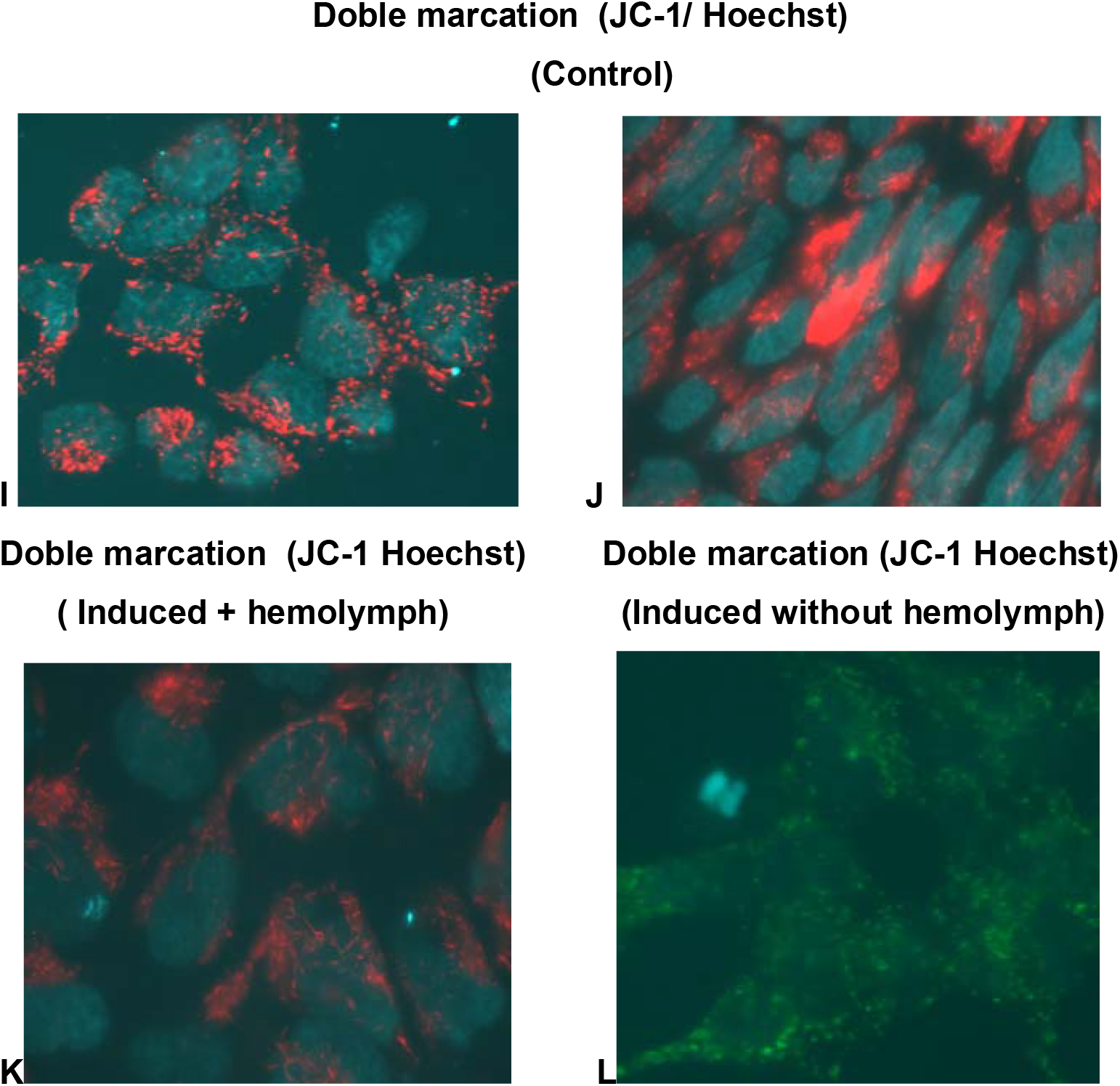
Photomicrography of HEK-293 cells without induction of death cell (A-B-E-F), or after induction (C-D-G-H) by chemical agents (t-BHP, 1st column or H_2_0_2_, 2nd column). After eighteen hour, cells were stained with JC-1(2μM) and Hoechst 33324 (2μM) for 20 minutes. The samples were observed in fluorescence microscope with an increase of 40x. In I-J is showed the sobreposicion of the two colors in normal culture (without induction). In figure K, the culture were apoptosis induced and after this treated with hemolymph (sobreposicion of the two colors). In figure L, the culture were apoptosis induced but without hemolymph treatment (sobreposicion of the two colors).

#### Identification of cytochrome C in HEK 293 cells after induction of apoptosis and the protective action of hemolymph

Cytochrome C is the first mitochondrial intermembrane protein released into the cytosol, with apoptogenic action. The literature relate that, together with dATP, cytochrome C is required for proteolytic activation of procaspase-3. The no release of cytochrome C in the cytosol in cultures treated with hemolymph is an indicator of the hemolymph action as an inhibitor of apoptosis cascade activation.

It is reported that the action of Bcl-2, a protein witch known for anti-apoptotic action, is exactly at mitochondrial membrane level. To determine the hemolymph effect in the protection of apoptosis induced by **t-BHP**, HEK 293 cells were grown in 12 wells microplate. After semi confluence, cells were treated with 1% (v / v) of purified hemolymph fraction (PRF) showing anti-apoptotic activity. After contact by one hour the cells were treated with 25, 50 or 75 μM **t-BHP**. The cultures were kept in CO_2_ incubator for 4 hours and after this period washed with PBS and fixed in paraformaldehyde (4%) and picric acid (0.18%) and incubated for 45 minutes. The cells were then washed with PBS and treated with 3% BSA and incubated for 30 minutes. The cells were again washed with PBS and labelled with monoclonal anti cytocromo C for 2 hours. The reaction was revealed with an anti mouse IgG alkaline phosphatase conjugate for 75 minutes. After this, was added 20 μM of Hoechst and the cell culture was observed in a fluorescence microscope (Leica DMRB microscope). The results are shown in **Figure 10**. Was observed that, as in control cultures (without death’s induction) as well in induced cultures, but treated with hemolymph, a high fluorescence in the mitochondria membrane. However, in cultures were the induction was performed, but without hemolymph treatment, this fluorescence was not observed. With the increase of the membrane porosity, and with consequent release of cytochrome C in citosol, the cytochrome C is distributed in a diffuse mode and cannot be clearly identified.

**Figure 10.**
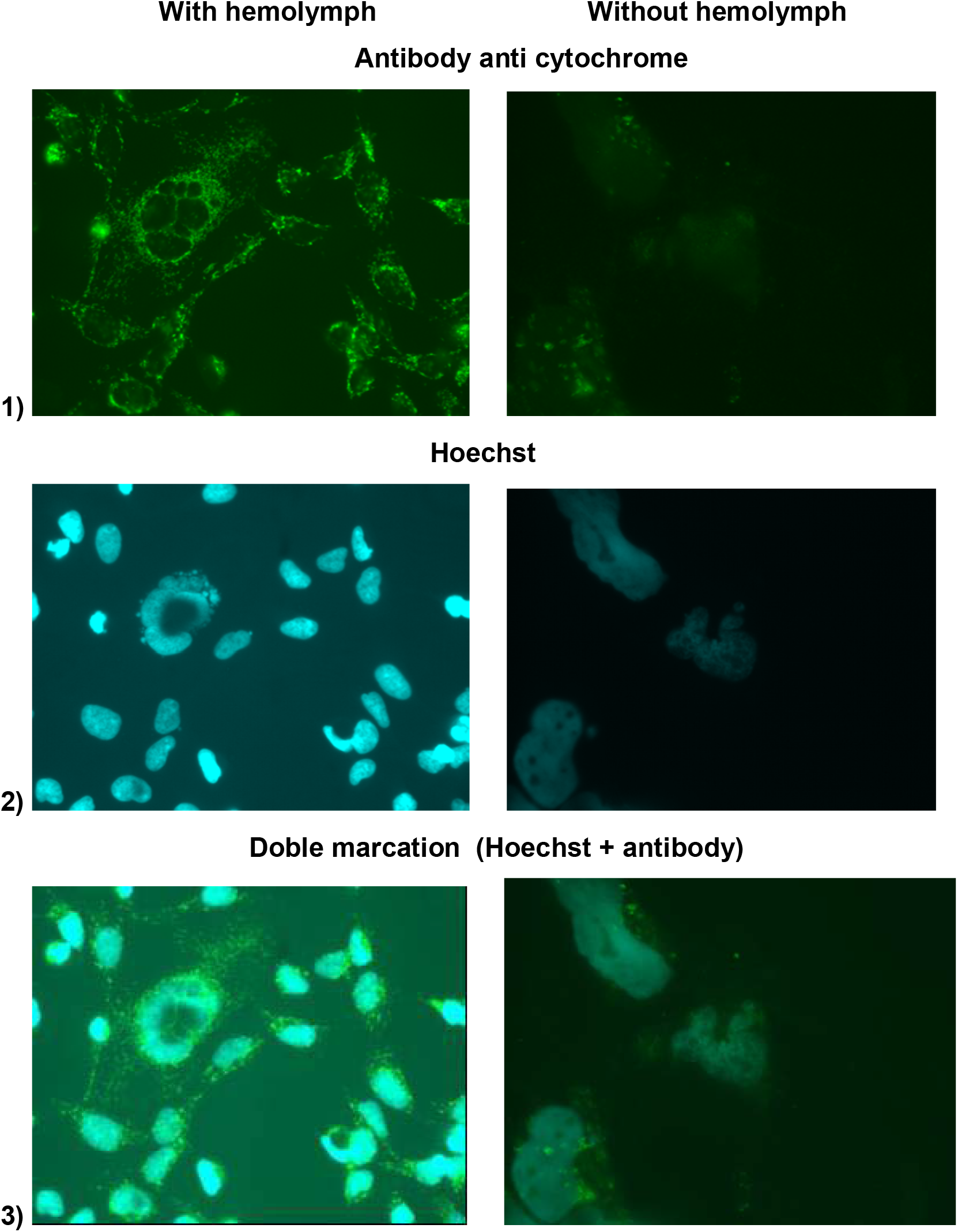
Photomicrography of HEK-293 cells cultured in microplate of 12 wells. After semi confluence the cells were treated with 1% (v/v) of the purified fraction of the hemolymph (Frp) showing antiapoptotic action. After one hour of contact the cells were treated with 25, 50 or 75μM of t-BPH. The cultures were maintained with CO_2_ by 4 hours and after this period washed with PBS and fastened with paraformaldeid 4% and picric acid 0,18% and incubated by 45 minutes. The cells were then stained with the antibody monoclonal anti cytochrome C 2. The reaction was revealed with an antibody anti mouse IgG conjugated with alkaline fosfatase by 75 minutes. Was added then 2μM of Hoechst and the cultures were observe in a fluorescence microscope with an increase of 40x. (A) cells treated with Frp (B) cells no treated with hemolymph. Both were induced with t-BHP

## DISCUSSION

Mitochondria are involved in many essential processes for cell survival, including energy production, redox control, calcium homeostasis and certain metabolic pathways and biosynthesis. In addition, the mitochondria have an essential role in the process of cell’s death known as apoptosis. This mitochondrial pathway of apoptosis may suffer dysfunctions causing various diseases such as cancer, diabetes, ischemia and neurodegenerative disorders like Parkinson’s and Alzheimer’s (Bouchier-Hayes et al. 2005). In one of the classic routes of apoptosis induction, the activation of caspases is directly related to mitochondrial outer membrane permeabilization (MOMP). Many signals that generate pro-apoptotic molecules and pathological stimulus converge on mitochondria and induce MOMP. The local regulation and for the execution of MOMP involve several types of proteins such as Bcl-2 family, mitochondrial lipids, proteins that regulate the metabolic flow and other components that act on the membranes pores. The MOMP is a lethal process because results in the release of capsize activating molecules and effectors of death that are caspase-independent. Some drugs that suppress the MOMP can prevents cells death or, when necessary, as in the case of cancer, restore the apoptosis process (Green and Kroemer 2005).

During the transduction of apoptosis signal, there is a change in the mitochondria membrane permeability, which causes the translocation of cytochrome C and apoptogenic proteins in the cytoplasm activatingb proteolytic proteins known as caspases. Shimizu et al (1999) has demonstrated that some pro-apoptotic proteins (Bax, Bak, etc.) would act on some voltage dependent of the anion channels (VDAC) accelerating the opening of these channels and increasing the permeability of the membrane, allowing that pro apoptotic activators factors of caspases proceed to the cytosol. On the other side, they also demonstrate that anti-apoptotic proteins family, known as Bcl-x (L) closes these channels by binding directly to them. While the proteins Bax and Bak allow cytochrome C to pass through these channels VDAC, keeping them open, the protein Bcl-x (L) prevents this occurring, regulating thus the mitochondrial membrane potential (ΔΨ_m_), blocking the release of cytochrome C during apoptosis-inducing events. On the other hand, Shimizu et al (2001) has shown that in some cases as the BH3 proteins (Bid and Bik) may have apoptogenic induction releasing cytochrome C by mechanisms other than those requiring the direct link to VDAC. Antibodies that inhibit VDAC prevent cytochrome release by the action of Bax avoiding the loss of mitochondrial membrane permeability. However, this is not observed with induction of release of cytochrome C by Bid. These anti-VDAC antibodies also inhibit apoptosis induced by chemical agents, inducers of apoptosis such as etoposide, paclitaxel and staurosporine. However, Cartier et al (1994) observed that, despite the fact that family of the proteins Bcl-2 are very efficient in blocking apoptosis in mammalian cells they fail to prevent death’s induced by actinomycin-D or induced by baculovirus (AcMNPV). Unlike what happens with the Bcl-2, p35, a potent inhibitor of apoptosis present in the genome of baculovirus, prevents apoptosis induced by this class of virus suggesting that the mechanism of action of two regulators may be functionally distinct. Due numerous mechanisms of the death apoptosis activation, the determination of the action mechanism of various anti-apoptotic proteins, particularly those of a broad spectrum of action (which inhibit death by different inducers or preventing this death in different types of cells) may be of high medical and biotechnological interest.

We have reported earlier the presence of anti-apoptotic protein present in the hemolymph of *Lonomia oblique* with his supplementation in cell’s culture inducing high levels of cell growth (Mendonça et al. 2008; Souza et al. 2005; Mendonça et al. 2002) and that the cells viability can be maintained for longer periods (Vieira et al. 2010; Souza et al. 2005; Maranga et al. 2003). However, contrary that occurs with the anti-apoptotic Bcl-2 family proteins (Shimizu et al. 1999), has the ability to inhibit apoptosis in different types of cells, whether mammal, either insect, by different inducers (Souza et al. 2005; Vieira et al. 2010). The present data suggests that the site of action of this protein could be similar to Bcl (x) proteins, since that, in all experiments, as we observed a blockade of apoptosis, with clear protection of mitochondrial membrane potential. The action of hemolymph on the cells is clearly demonstrated in early experiments, whether in the cell viability increase, either in the increased of the recombinant proteins production (Vieira et al. 2010). This action, at least in part related to an anti-apoptotic protein, which is evident in with the cell’s death blocking, the cell remains functional **Figures 4–8**. In this case we have observed increasing the viral title or in the recombinant proteins production (Mendonça et al. 2008). As demonstrated throughout the study, apparently the site of action of this protein could be on the membrane of mitochondria, maintaining so, the high membrane potential, avoiding the membrane permeability. This anti-apoptotic protein not only blocks death induced by the virus, but also death’s that normally occurs by physiological stress culture, as observed in control culture ( **Figure 8**). In this case, the cell’s death observed in the control culture was reduced by half, indicating that this protein could also be used as a supplement to culture media. The protein responsible for this effect has been identified as a protein with a molecular weight of 51 kDa (Souza et al. 2005). The protective effect of p51 protein is evident in **Figures 4–8**, where death was induced with a potent cell death inducer (t-BHP or H_2_0_2_). Under these conditions p51 was able to inhibit the death induced in insect cells (Sf-9) at concentrations above 50 μM t-BHP or 400 μM of hydrogen peroxide.

Flow cytometry proved an excellent tool to characterize different cell populations. This technique was used in all experiments used to determination of potential of the mitochondrial membrane as well apoptosis cell’s death (figures not showed). The definition of the different cells population (viable, necrotic and apoptotic cells) were determined by information obtained from literature. Apoptotic cells show small size (low FSC) and high granularity (high SSC) and intensely are stained by PI. These cells showed too low potential of the mitochondrial membrane. Necrotic cells are stained by PI. The differentiation between necrotic from apoptotic death is that in necrotic cells shown a normal size with a high potential of the mitochondrial membrane. Normal cells show a high viability, high potential of the mitochondrial membrane and are not labelled with PI. Was observed a material showing low size and low granularity and are not stained by no markers. This material are cellular debris or apoptotic bodies. The characteristic of each of these populations has been confirmed by other techniques such as electron microscopy, and agarosis gel electrophoresis to determine DNA fragmentation (Petit et al. 1995; Vieira et al. 2002). The determination of DIOC6 (3) was effective potential of the mitochondrial membrane, determination. However, to confirm that the hemolymph action site is really on the mitochondria, we used a second dye (JC-1). This stain been widely used by other researchers for this purpose. The results obtained with this marker were similar to those obtained with the DIOC6 (3). Also was observed, in all the experiments, cell death in the control culture (without induction) when the incubation is performed overnight.

Nevertheless, when hemolymph was added to cell culture it was able to inhibit virtually 100% of the harmful effect observed in culture.

The cytochome C is the first protein intermembrana mitochondrial liberated in the cytosol, with action apoptogenic. It is mentioned in the literature that, together with the dATP, the cytochrome C is necessary for the proteolític activation of the procaspase-3. If the cytochrome C is not liberated in the cytosol in the cell’s culture treated with hemolymph, it would be an indicator of the hemolymph actions in the inhibition of this cascade need to apoptosis activation. According Zou et al., (1997), the cytochrome C attached to Apaf-1, which in presence of ATP (or dATP), promote the activation of the procaspase-9, generating the activated caspase-9 (Li et al. 1997). Therefore, caspase-9 act on the procaspase-3 to begin the activation of the caspase cascade. If the hemolymph were acting about the liberation of the cytochrome C in the cytosol, as we suspected, there would be a suppression of the activation of the caspase-3, blocking this way the apoptosis induction. This blocked of the cytochrome C maintains the functionality of the mitochondria, avoiding the lack of energy for the cell, and therefore the cellular death by apoptosis. In this study, we demonstrate the action of hemolymph on the mitochondria and release of cytochrome C in cytosol. As expected, when cultures were treated with hemolymph, there was no release of cytochrome C, which remained fixed on the membrane of mitochondria, generating an intense fluorescence. Meanwhile, in cultures induced with t-BHP and non-treated hemolymph, this was not observed.

## Abbreviations

Hb: hemolymph
t-BHP: *tert*-butylhydroperoxide
PI: propidium iodide
DiOC_6_(3): 3,3’dihexyloxacarbocyanine iodide
JC-1,3: 3’e-tetraethylbenzimidazolylcar-bocyanine iodide
hpi: hours post infection
MOI: multiplicity of infection
MMP: mitochondrial membrane permeabilization

## Acknowledgements

The authors acknowledge the financial support of FAPESP (2009/06511-1) and the technical assistance of Ana C.P. Pereira, Helena L.A. Vieira and Cristina C. Peixoto from Instituto de Biologia Experimental e Tecnológica (IBET), Oeiras, Portugal

## REFERENCE

Arden N, Ahn SH, Vaz W, Rhodes M, Hancock C, Abitorabi MA, Betenbaugh MJ (2007) Chemical caspase inhibitors enhance cell culture viabilities and protein titer. Biotechnol Prog 23 (2):506–511.

Bedoui, S., Herold, M. J., and Strasser, A. (2020). Emerging connectivity of programmed cell death pathways and its physiological implications. Nat. Rev. Mol. Cell Biol. 21, 678–695

Bock, F. J., and Tait, S. W. G. (2020). Mitochondria as multifaceted regulators of cell death. Nat. Rev. Mol. Cell Biol. 21, 85–100

Bouchier-Hayes L, Lartigue L, Newmeyer DD (2005) Mitochondria: pharmacological manipulation of cell death. J Clin Invest 115 (10):2640–2647.

Bradford MM (1976) A rapid and sensitive method for the quantitation of microgram quantities of protein utilizing the principle of protein-dye binding. Anal Biochem 72:248–254.

Butler M (2005) Animal cell cultures: recent achievements and perspectives in the production of biopharmaceuticals. Appl Microbiol Biotechnol 68 (3):283–291

Cao, W., Li, J., Yang, K., and Cao, D. (2021). An overview of autophagy: mechanism, regulation and research progress. Bull. Cancer 108, 304–322

Cartier JL, Hershberger PA, Friesen PD (1994) Suppression of apoptosis in insect cells stably transfected with baculovirus p35: dominant interference by N-terminal sequences p35(1-76). J Virol 68 (12):7728–7737

Ferri KF, Jacotot E, Blanco J, Este JA, Zamzami N, Susin SA, Xie Z, Brothers G, Reed JC, Penninger JM, Kroemer G (2000) Apoptosis control in syncytia induced by the HIV type 1-envelope glycoprotein complex: role of mitochondria and caspases. J Exp Med 192 (8):1081–1092

Ferri KF, Kroemer G (2001) Organelle-specific initiation of cell death pathways. Nat Cell Biol 3 (11):E255–263.

Green DR, Kroemer G (2005) Pharmacological manipulation of cell death: clinical applications in sight? J Clin Invest 115 (10):2610–2617.

Kazek, M., Kaczmarek, A., Wronska, A. K., and Bogus, M. I. (2020). Conidiobolus coronatus induces oxidative stress and autophagy response in *Galleria mellonella* larvae. PLoS One 15:e0228407.

Kesavardhana, S., Malireddi, R. K. S., and Kanneganti, T. D. (2020). Caspases in cell death, inflammation, and pyroptosis. Annu. Rev. Immunol. 38, 567–595. doi: 10.1146/annurev-immunol-073119-095439

Kroemer G, Galluzzi L, Brenner C (2007) Mitochondrial membrane permeabilization in cell death. Physiol Rev 87 (1):99–163.

Li P, Nijhawan D, Budihardjo I, Srinivasula SM, Ahmad M, Alnemri ES, Wang X (1997) Cytochrome c and dATP-dependent formation of Apaf-1/caspase-9 complex initiates an apoptotic protease cascade. Cell 91 (4):479–489.

Maranga L, Mendonca RZ, Bengala A, Peixoto CC, Moraes RH, Pereira CA, Carrondo MJ (2003) Enhancement of Sf-9 cell growth and longevity through supplementation of culture medium with hemolymph. Biotechnol Prog 19 (1):58–63.

Mastrangelo AJ, Zou S, Hardwick JM, Betenbaugh MJ (1999) Antiapoptosis chemicals prolong productive lifetimes of mammalian cells upon Sindbis virus vector infection. Biotechnol Bioeng 65 (3):298–305.

Mendonça RZ, Arrozio SJ, Antoniazzi MM, Ferreira JM, Jr., Pereira CA (2002) Metabolic active-high density VERO cell cultures on microcarriers following apoptosis prevention by galactose/glutamine feeding. J Biotechnol 97 (1):13–22.

Mendonça RZ, Greco KN, Sousa AP, Moraes RH, Astray RM, Pereira CA (2008) Enhancing effect of a protein from *Lonomia obliqua* hemolymph on recombinant protein production. Cytotechnology 57 (1):83–91.

Meneses-Acosta A, Mendonça R, Merchant H, Covarrubias L, Ramirez O (2001) Comparative characterization of cell death between Sf9 insect cells and hybridoma cultures. Biotechnol Bioeng 72 (4):441–457

Nury, T., Zarrouk, A., Yammine, A., Mackrill, J. J., Vejux, A., and Lizard, G. (2021). Oxiapoptophagy: a type of cell death induced by some oxysterols. Br. J. Pharmacol. 178, 3115–3123.

Petit PX, Lecoeur H, Zorn E, Dauguet C, Mignotte B, Gougeon ML (1995) Alterations in mitochondrial structure and function are early events of dexamethasone-induced thymocyte apoptosis. J Cell Biol 130 (1):157–167

Sauerwald TM, Oyler GA, Betenbaugh MJ (2003) Study of caspase inhibitors for limiting death in mammalian cell culture. Biotechnol Bioeng 81 (3):329–340.

Shimizu S, Matsuoka Y, Shinohara Y, Yoneda Y, Tsujimoto Y (2001) Essential role of voltage-dependent anion channel in various forms of apoptosis in mammalian cells. J Cell Biol 152 (2):237–250

Shimizu S, Narita M, Tsujimoto Y (1999) Bcl-2 family proteins regulate the release of apoptogenic cytochrome c by the mitochondrial channel VDAC. Nature 399 (6735):483–487

Souza AP, Peixoto CC, Maranga L, Carvalhal AV, Moraes RH, Mendonça RM, Pereira CA, Carrondo MJ, Mendonca RZ (2005) Purification and characterization of an anti-apoptotic protein isolated from *Lonomia obliqua* hemolymph. Biotechnol Prog 21 (1):99–105.

Souza AP, Moraes RH, Mendonca RZ (2015) Improvement replication of the Baculovirus Anticarsia gemmatalis nucleopolyhedrovirus (AgMNPV) in vitro using proteins from Lonomia oblique hemolymph. Cytotechnology, (2015) 67 (2):331–34.2

Vieira HL, Boya P, Cohen I, El Hamel C, Haouzi D, Druillenec S, Belzacq AS, Brenner C, Roques B, Kroemer G (2002) Cell permeable BH3-peptides overcome the cytoprotective effect of Bcl-2 and Bcl-X(L). Oncogene 21 (13):1963–1977.

Vieira HL, Pereira AC, Carrondo MJ, Alves PM (2006) Catalase effect on cell death for the improvement of recombinant protein production in baculovirus-insect cell system. Bioprocess Biosyst Eng 29 (5-6):409–414.

Vieira HL, Pereira AC, Peixoto CC, Moraes RH, Alves PM, Mendonça RZ (2010) Improvement of recombinant protein production by an anti-apoptotic protein from hemolymph of *Lonomia obliqua*. Cytotechnology 62 (6):547–555.

Zou H, Henzel WJ, Liu X, Lutschg A, Wang X (1997) Apaf-1, a human protein homologous to C. elegans CED-4, participates in cytochrome c-dependent activation of caspase-3. Cell 90 (3):405–413.

Wang, Y., Goodman, C. L., Ringbauer, J. Jr., Li, Y., and Stanley, D. (2021). Prostaglandin A2 induces apoptosis in three cell lines derived from the fall armyworm, *Spodoptera frugiperda*. Arch. Insect Biochem. Physiol. 108:e21844.

Wrońska A.K, Kaczmarek A, Kazek M and Bogun M.I. Infection of *Galleria mellonella* (Lepidoptera) Larvae With the Entomopathogenic Fungus *Conidiobolus coronatus* (Entomophthorales) Induces Apoptosis of Hemocytes and Affects the Concentration of Eicosanoids in the Hemolymph Front. Physiol., 06 January 2022

